# leADS: improved metabolic pathway inference based on active dataset subsampling

**DOI:** 10.1101/2020.09.14.297424

**Authors:** Abdur Rahman M. A. Basher, Aditi N. Nallan, Ryan J. McLaughlin, Julia Anstett, Steven J. Hallam

**Affiliations:** Graduate Program in Bioinformatics, University of British Columbia, Genome Sciences Centre, 100-570 West 7th Avenue, Vancouver, British Columbia V5Z 4S6, Canada; Department of Microbiology & Immunology, University of British Columbia, 2552-2350 Health Sciences Mall, Vancouver, British Columbia V6T 1Z3, Canada; Genome Science and Technology Program, University of British Columbia, 2329 West Mall, Vancouver, BC V6T 1Z4, Canada; Life Sciences Institute, University of British Columbia, Vancouver, British Columbia, Canada V6T 1Z3; ECOSCOPE Training Program, University of British Columbia, Vancouver, British Columbia, Canada V6T 1Z3

## Abstract

Metabolic pathways are composed of reaction sequences catalyzed by enzymes. The set of reactions within and between cells comprises a reactome. Pathways and reactomes can be predicted from organismal or multi-organismal genomes using rule-based or machine learning methods. While machine learning methods overcome issues of probability and scale associated with rule-based methods, several complications remain that can degrade performance including inadequately labeled training data, missing feature information, and inherent imbalances in the distribution of pathways within a dataset. Here, we present leADS (mu<u>l</u>ti-label l<u>e</u>arning based on <u>a</u>ctive <u>d</u>ataset <u>s</u>ubsampling), a machine learning method, that uses subsampling to reduce the negative impact of training loss due to class imbalance. We demonstrate leADs performance using organismal and multi-organismal datasets in relation to other machine learning pathway prediction methods.

**Availability and implementation:** leADS is available under the GNU license at github.com/hallamlab/leADS. A wiki, including a tutorial, is available at github.com//hallamlab/leADS/wiki

## 1 Introduction

The rise of next generation sequencing technologies has motivated innovations in metabolic pathway prediction methods [1]. These innovations encompass rule-based or heuristic methods including PathoLogic [14], and machine learning (ML) methods including PtwML [6] and mlLGPR [20]. In the ML case, a class imbalance problems exists where certain pathways are more common than others because they conduct core metabolic functions conserved across the tree of life. These functions are overrepresented in labeled training data relative to more niche-defining or accessory metabolic functions and can result in training loss with decreased predictive performance. To address this problem, we developed leADS based on prior work in dataset subsampling [10]. leADS incorporates an ensemble of multi-label learners [32] to perform hard example mining [27], reducing the negative impact of training loss on pathway prediction.

## 2 Methods

leADS (Fig. 1A.) performs training in three iterative steps:

**Figure 1:**
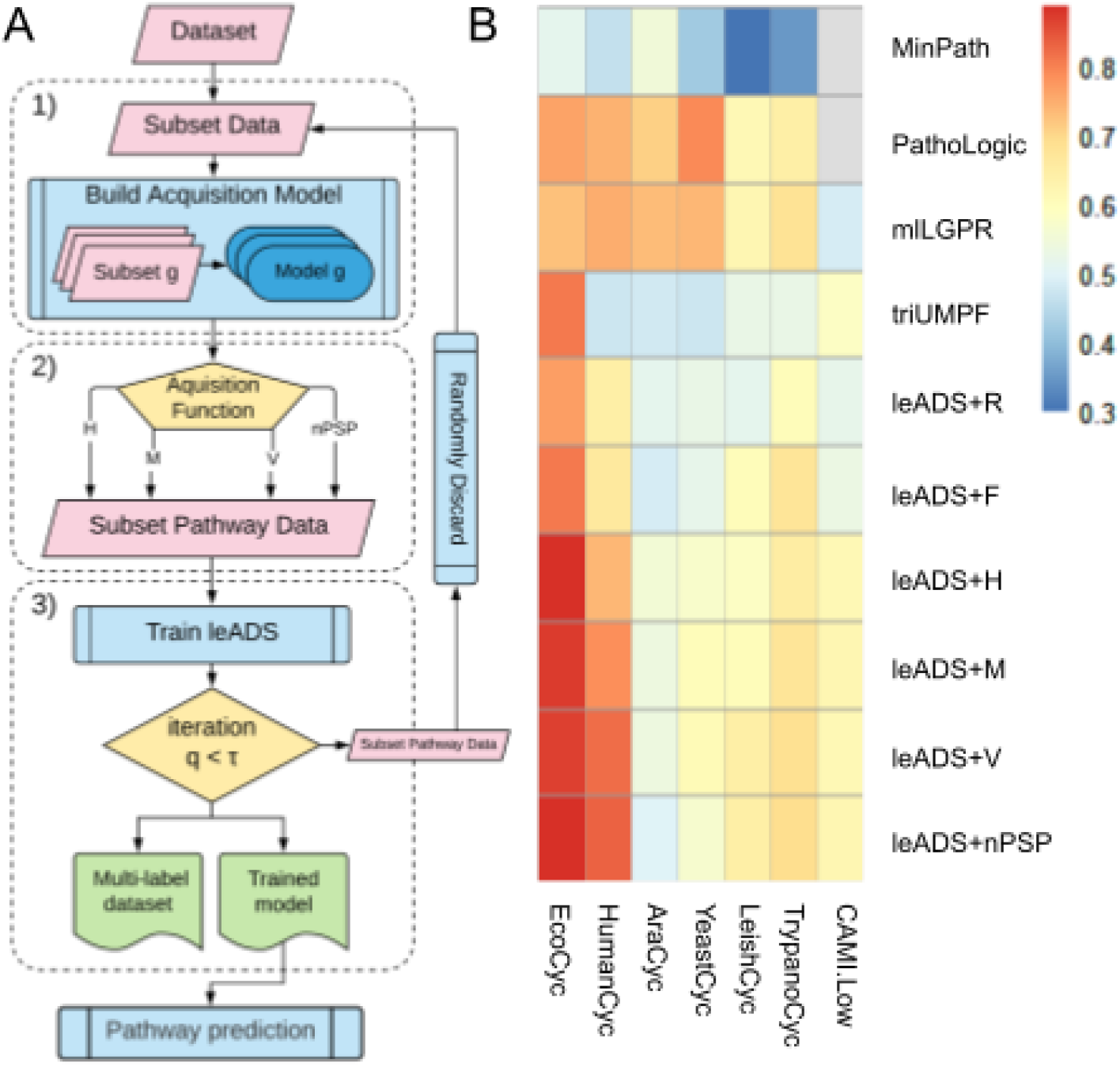
Fig. A)- leADS workflow. Fig B)- predictive performance of MinPath, Pathologic, mlLGPR, triUMPF, and leADS (with random sampling, full data, and four acquisition functions) on Tier 1 (T1) organismal genomes and Critical Assessment of Metagenome Interpretation (CAMI) datasets. (x-axis). Z-axis is average F1 score. Gray cells indicate that algorithm was not run on that dataset.

### 1)- Building an acquisition model

At the very first iteration, an empty set is initialized with randomly selected data from a given pathway dataset (Fig. 1A.1). Then, an ensemble consisting of *g*(∈𝕫_≥1_) members is constructed, where each member *g* in the ensemble is trained on a randomly selected subset of the data.

### 2)- Dataset sub-sampling

During this step, a subset of pathway data is selected using one of four acquisition functions including *entropy (H), mutual information (M), variation ratios (V)*, or *normalized propensity scored precision at k* ∈𝕫_*>*1_ *(nPSP)* (Fig. 1A.2). For each function, the top **per**% examples are retrieved, where **per**%(∈(0, 100]) is a prespecified hyperparameter indicating the subsampling proportion.

### 3)- Train using sub-sampled data

The selected subset of pathway data from the previous step are used to train leADS using a multi-label 1-vs-All approach [20] (Fig. 1A.3).These steps are repeated *τ* (∈𝕫_≥1_) times (Fig. 1A). For each iteration some examples collected from the previous iteration *q* −1 are randomly discarded to enable examples not selected in the top **per**% to be used in round *q*. Once training is complete: i)- pathway data with **per**% examples is produced; and ii)- the trained model is stored to use in pathway prediction on new datasets. For definitions, analytical expressions, and optimization, consult Supp. Sections A.1, A.2, and A.3.

We evaluated leADS performance using a corpora of 10 experimental datasets manifesting diverse multi-label properties, including manually curated organismal genomes, synthetic microbial communities and low complexity microbial communities. We trained leADS on BioCyc v21 Tier 2 and Tier 3 (T2 &3) under three configurations: i)- random sampling (leADS+R) corresponding to 70% of BioCyc v21 T2 &3 selected at random, ii)- full configuration (leADS+F) where all BioCyc v21 T2 &3 data were utilized without subsampling, and iii)- **per**% = 70% using four acquisition functions: entropy (leADS+H), mutual information (leADS+M), variation ratios (leADS+V), and normalized propensity scored precision (leADS+nPSP). Training for each configuration was run for 10 epochs using *g* = 10 member size and *k* = 50 (for leADS+V and leADS+nPSP). For detailed experimental settings, see Supp. Section A.5.3. Pathway prediction results are reported using the *average F1 score*. As shown in Fig. 1B, leADS resulted in competitive performance compared to other inference methods. Among the four acquisition functions, nPSP resulted in the highest performance on EcoCyc (0.8874) while random sampling resulted in the poorest. Interestingly, full was on par with random sampling, reinforcing the idea that BioCyc T2 &3 contains noise that may hamper proper estimation of leADS coefficients. On CAMI low complexity metagenomes ([24]), nPSP outperformed other methods (0.6214) Fig. 1B. Based on these results we recommend using nPSP with *g* = 10 and *k* = 50 settings for optimal leADS performance. Extensive experimental analysis can be found in Supp. Sections A.4 and A.5.

## 3 Conclusion

leADS is a novel multi-label ensemble-based approach for hard example mining that constructs a set of diverse multi-label base learners to jointly improve subselection of examples and overcome class imbalance during metabolic pathway prediction from genomic sequence information at different levels of complexity and completion.

## A Supplementary Material

This material is divided into five parts: i)- the problem definitions (Section A.1), ii)- the leADS framework (Section A.2), iii)- optimization and prediction (Section A.3), iv)- experimental settings (Section A.4), and v)- empirical analysis (parameter sensitivity, scalability to the ensemble size, and metabolic pathway prediction effectiveness) (Section A.5).

### A.1 Definitions and Problem Formulation

Here the default vector is considered to be a column vector and is represented by a boldface lowercase letter (e.g., **x**) while the matrices are represented by boldface uppercase letters (e.g., **X**). If a subscript letter *i* is attached to a matrix, such as **X**_*i*_, it indicates the *i*-th row of **X**, which is a row vector. A subscript character to a vector, **x**_*i*_, denotes an *i*-th cell of **x**. Occasional superscript, **x**^(*i*)^, suggests an index to an example or current epoch during a learning period. With these notations in mind, we introduce information integral to the problem formulation, starting by defining the multi-label data.

#### Definition A.1.

Multi-label Pathway Dataset [20]. A pathway dataset is represented by *𝒮*_*A*_ = {(**x**^(*i*)^, **y**^(*i*)^) : 1 *< i ⩽ n*} consisting of *n* examples, where **x**^(*i*)^ is a vector indicating the abundance information corresponding to enzymatic reactions. An enzymatic reaction is denoted by *c*, which is an element of a set ε= {*c*_1_, *c*_2_, …, *c*_*r*_}, having *r* possible enzymatic reactions, hence, the vector size **x**^(*i*)^ is *r*. The abundance of an enzymatic reaction for an example *i*, say 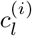, is defined as 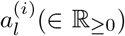. The class labels 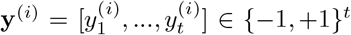 is a pathway label vector of size *t* representing the total 𝓎 number of pathways derived from a set of universal metabolic pathway. The matrix form of **x**^(*i*)^ and **y**^(*i*)^ are **X** and **Y**, respectively. ▪

Both ε and 𝓎 can be retrieved from trusted sources, such as KEGG [12] or MetaCyc [3]. Although the input space is assumed to be encoded as *r*-dimensional vector, symbolized as 𝒳 = ℝ^*r*^, through features engineering it can be represented as X = ℝ^*d*^.

#### Problem Statement 1.

*Given a multi-label dataset*, 𝒮_*A*_, *the goal is to select a subset of* 𝒮_*A*_, *denoted by* 𝒮_***per***%_, *where* **per**% *is a prespecified hyperparameter, indicating the proportion of examples to be chosen from* 𝒮_*A*_, *such that learning on* 𝒮_***per***%_ *incurs similar predictive score (or better) as if it was trained on full multi-label dataset*, 𝒮_*A*_.

### A.2 The leADS Method

In this section, we provide a description of leADS components including: i)- building an acquisition model, ii)- active dataset sub-sampling, and iii)- learning using the reduced sub-sampled data. These three steps interact with each other in an iterative process as illustrated in Fig. 2. At the very first iteration, a set 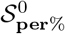 is initialized with randomly selected data (Fig. 2a-b). In the next iteration *q*, instead of re-initializing 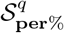 with randomly selected examples, 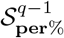 data collected from the previous iteration *q* −1 is used, constituting a *build-up scheme* implemented in many active learning methods [5, 10]. This process is repeated until the maximum number of rounds *τ* is reached.

**Figure 2:**
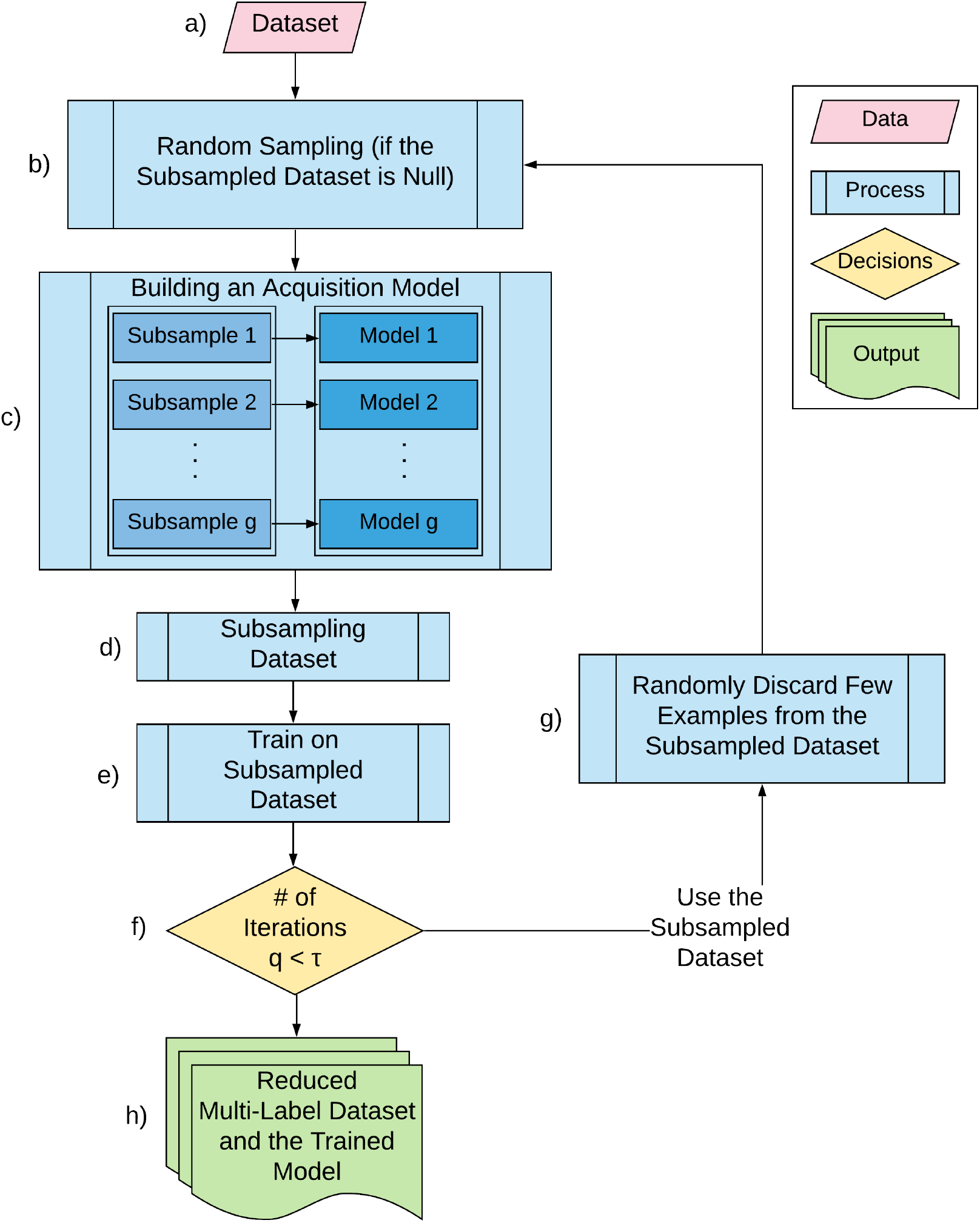
A schematic diagram indicating the leADS workflow. Using a multi-label pathway dataset (a), leADS randomly selects examples at the very first iteration (b) then builds *g* members of an ensemble (c), where each is trained on a randomly selected portion of the training set. Next, leADS applies an acquisition function (d), based on either: entropy, mutual information, variation ratios, or normalized propensity scored precision at *k*, to select **per**% sub-samples. Then, leADS performs parallel training steps (e). The process (b-e) is repeated *τ* times (f), where during each iteration few examples from **per**% are discarded at random (g) to give chance to examples that were not selected in **per**% to be used for the next subsequent round for training. If the current iteration *q* reaches a desired number of rounds *τ*, training is terminated and the final model and **per**% results are presented (h).

#### A.2.1 Building an Acquisition Model

Given 𝒮_*A*_, the objective of this step (Fig. 2c) is to estimate posterior predictive uncertainty given a new test example **x**^∗^ for a pathway **y**_*j*_ as:

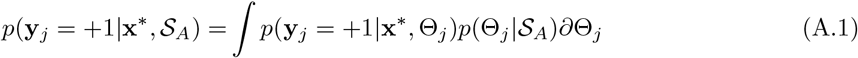

where Θ ∈ ℝ^*t*×*r*^ denotes pathway’s parameters. Notice that Eq A.1 involves marginalization over Θ_*j*_ parameters, which is hard to compute [22]. One way to mitigate this issue is to approximate the above equation using Monte Carlo (MC) techniques [16] by constructing an ensemble, denoted by *E*, which consists of *g*(∈𝕫_≥1_) models (Fig. 2c) where each generates multiple examples according to the following formula:

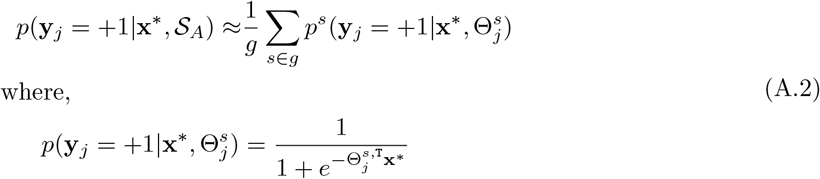

where 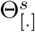 is sampled from *q*(Θ^*s*^) which is considered to be in the same family distribution as the true hidden variables 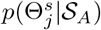. The parameters Θ^*s*^ for the *s*-th model can be estimated according to the multi-label 1-vs-All approach [32].

#### A.2.2 Subsampling Dataset

During this step (Fig. 2d), a subset of 𝒮_*A*_, denoted as 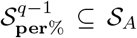, is picked for each member in *E* using an acquisition function *f* : **x** → ℝ where **per**% is a pre-specified threshold, indicating the proportion of examples to be chosen from 𝒮_*A*_, at iteration *q* −1.

Four acquisition functions used in subsampling are described that incorporate predictive uncertainty distribution from the previous step: *entropy, mutual information, variation ratios*, and *normalized PSP*@*k*. For each function, we retrieve top **per**% examples that contain high acquisition (or uncertainty) values.

1. **Entropy (ℋ)** [25]. This function measures the uncertainty of an example given the predictive distribution of that example:

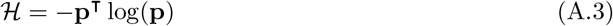

where **p** is a vector of predictive probabilities over *t* pathways.
2. **Mutual information (ℳ)** [28]. This function looks for low mutual information between *g* models, encouraging examples with high disagreement to be selected during the data acquisition process:

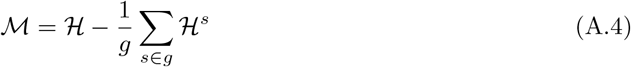

where ℋ ^*s*^ denotes the entropy obtained from an individual member of *E* for an example before marginalization. Since entropy is always positive, the maximum possible value for ℳ is ℋ. However, when the models make similar predictions, then 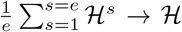, resulting in ℳ → 0, which is its minimum value ([5]). Note that this formula is similar to multi-label negative correlation learning ([26]), which estimates pairwise negative correlation of each learner’s error with respect to errors of other members in *E*.
3. **Variation ratios** (𝒱) [7]. This function measures the number of members in *E* that disagree with the majority vote for an example according to *k* desired pathway size, where larger values indicate higher uncertainty:

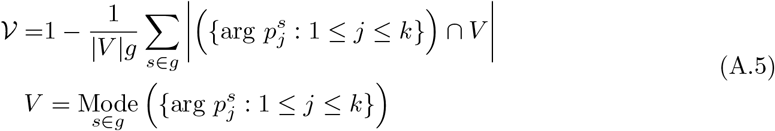

where *V* corresponds the disagreement of *k* pathways across *g* models, where *k∈* 𝕫_*>*0_ is a prespecified number of pathways to be considered in computing the mode operation.
4. **Normalized propensity scored precision at** *k* (nPSP@*k*). This is a modified version of PSP@*k* [11], *which measures the average precision of top k* relevant pathways given an instance *i* where larger values indicate less uncertainty:

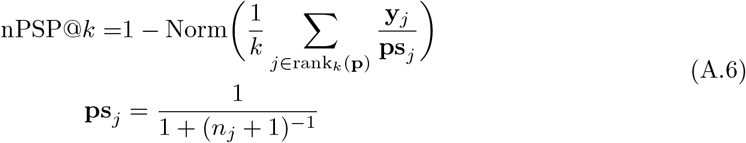

where Norm(.) scales the score within [0, 1]. The term **p** is a vector of predictive probabilities over *t* pathways, rank_*k*_(**p**) returns the indices of *k* largest value in **p**, ranked in a descending order, where *k* ∈ 𝕫_*>*0_ is a hyperparameter. **ps**_*j*_ is the propensity score for the *j*-th pathway, where *n*_*j*_ is the number of the positive training instances for the pathway *j*. In the context of extreme multi-label problems, PSP@*k* was used to derive an upper bound for missing/miss-classified labels [30], and is reported to be the best suited metric for long-tail distribution in which a significant portion of labels are tail labels [23, 2].

#### A.2.3 Train on the Reduced Dataset

During this step (Fig. 2e), each member in *E* is assigned to train on randomly selected examples from 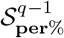, which is expected to contain hard examples that are difficult to learn and classify. This process is repeated *τ* times (Fig. 2f), where during each iteration the top **per**% are selected based on their acquisition values for the next round of training. However, leADS discard few examples (*γ* ∈ (0, 1)) from 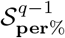 to increase coverage of information from all examples for the next round (Fig. 2g).

### A.3 Optimization and Prediction

The objective function in Eq. A.2 can be solved by decomposing into *t* independent binary classification problems according to the multi-label 1-vs-All approach enabling parallel training. Consider optimization for a member *s*:

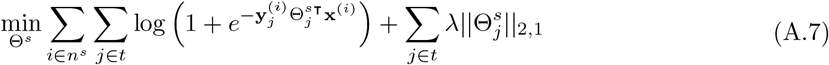

where 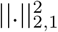 is the L_2,1_ regularization term, which is the sum of the Euclidean norms of columns of Θ. The L_2,1_ norm imposes sparsity on the model’s parameters to minimize the negative effect of label correlations, where *λ*(∈ℝ_*>*0_) is employed to govern relative contributions of L_2,1_ and the log-loss term. Although the joint formula in Eq A.7 is convex, the logistic log-loss function still posses a problem where there exists no analytical solution for it. To address this problem, we apply mini-batch gradient descent [18], which begins with some initial random guess for leADS parameters, and performs iterative updates to each individual parameter to minimize Eq. A.7 where the derivative for each 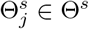 has the following formula:

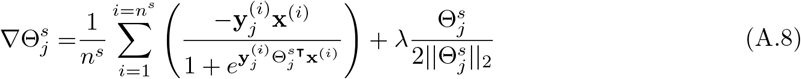

For prediction, we apply a cut-off threshold *ξ* ∈ ℝ_≥0_ to retain only pathways having higher probability values than *ξ*, i.e., 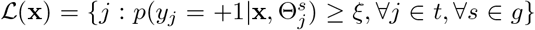, where 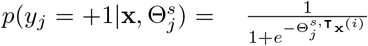.

### A.4 Experimental Setup

In this section, we describe an experimental framework used to demonstrate leADS pathway prediction performance across multiple datasets spanning the genomic information hierarchy [20]. leADS was written in the Python programming language (v3). Unless otherwise specified all tests were conducted on a Linux server using 10 cores of Intel Xeon CPU E5-2650.

#### A.4.1 Description of Datasets

We used a corpora of 10 organismal and multi-organismal datasets including BioCyc v21 T2 & 3 [4], T1 PGDBs from the BioCyc collection (*EcoCyc (v21), HumanCyc (v19*.*5), AraCyc (v18*.*5), YeastCyc (v19*.*5), LeishCyc (v19*.*5)*, and *TrypanoCyc (v18*.*5)*), symbiont genomes describing distributed metabolic pathways for 9 amino acid biosynthesis between the two symbiotic bacteria: *Moranella* (GenBank NC-015735) living inside *Tremblaya* (GenBank NC-015736) [21], genomes used in the CAMI initiative [24], and whole genome shotgun sequences from HOTS at 25m, 75m, 110m (sunlit) and 500m (dark) ocean depth intervals [29], to benchmark leADS. MetaCyc database version 21 [3] (composed of 2526 base pathways and 3650 enzymatic reactions) was used as a trusted source to refine these 𝒮datasets by including only those pathways that intersect with this version of MetaCyc. The preprocessed experimental datasets can be obtained from https://doi.org/10.6084/m9.figshare.16752685. Table 1 summarizes the general characteristics of the applied datasets. For each dataset 𝒮, we use|𝒮 | and L(𝒮) to represent the number of instances and pathway labels, respectively. In addition, we also present some characteristics of the multi-label datasets, which are denoted as:

**Table 1:**
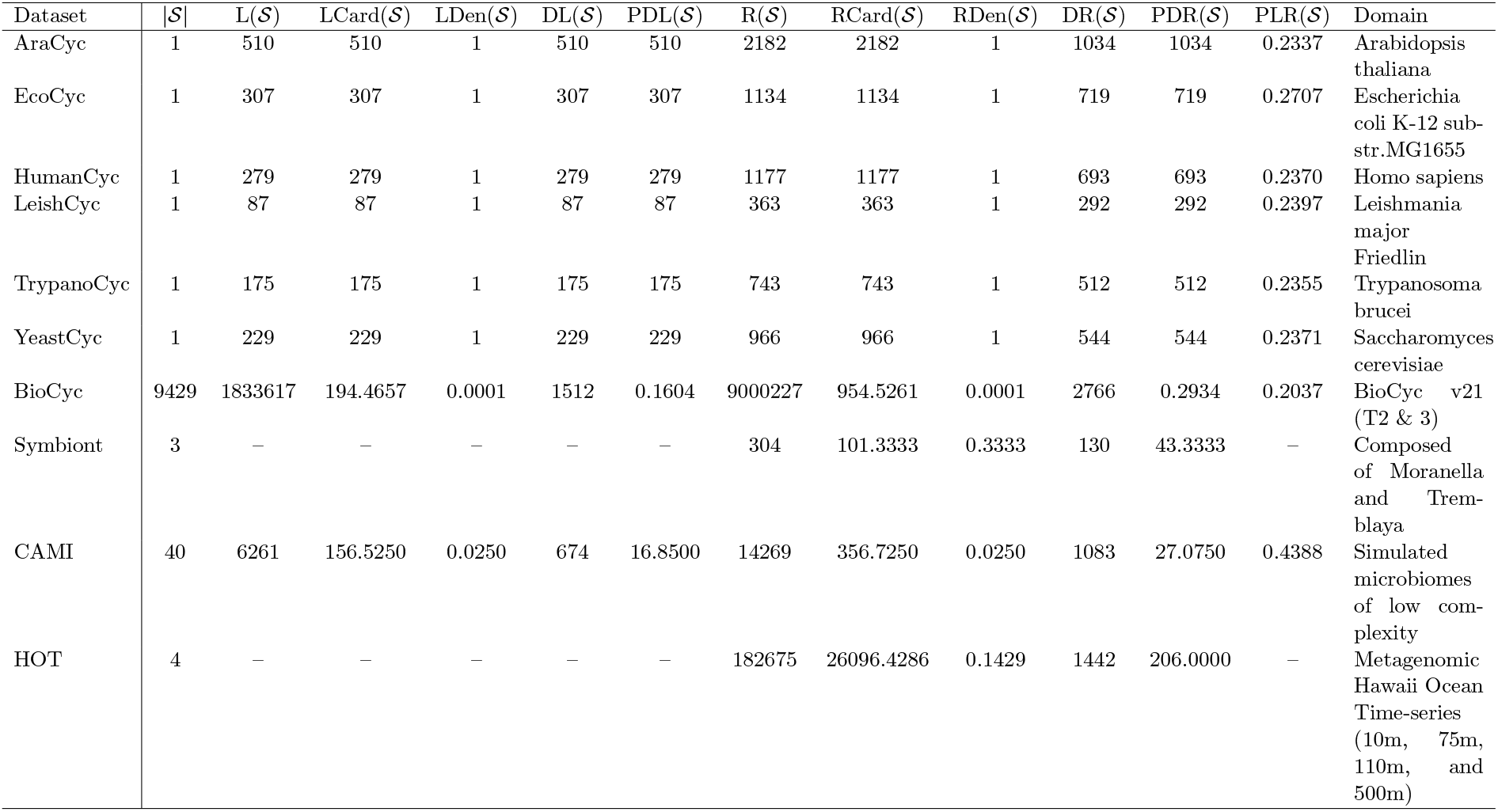
Experimental data set properties. The notations |𝒮|, L(𝒮), LCard(𝒮), LDen(𝒮), DL(𝒮), and PDL(𝒮) represent: number of instances, number of pathway labels, pathway labels cardinality, pathway labels density, distinct pathway labels, and proportion of distinct pathway labels for 𝒮, respectively. The notations R(𝒮), RCard(𝒮), RDen(𝒮), DR(𝒮), and PDR(𝒮) have similar meanings for the enzymatic reactions ε in S. PLR(𝒮)represents a ratio of L(𝒮) to R(𝒮). The last column denotes the domain of 𝒮.

1. Label cardinality 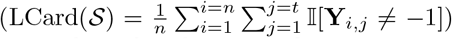, where 𝕀 is an indicator function. It denotes the average number of pathways in 𝒮
2. Label density 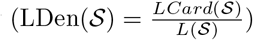. This is simply obtained through normalizing LCard(𝒮) by the number of total pathways in 𝒮.
3. Distinct pathway labels (DL(𝒮)). This notation indicates the number of distinct pathways in𝒮.
4. Proportion of distinct pathway labels 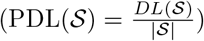. It represents the normalized version of DL(𝒮), and is obtained by dividing DL(𝒮.) with the number of instances in 𝒮.

The notations R(𝒮), RCard(𝒮), RDen(𝒮), DR(𝒮), and PDR(𝒮) have similar meanings for the enzymatic reactions ε in 𝒮. Finally, PLR(𝒮) represents a ratio of L(𝒮) to R(𝒮).

#### A.4.2 Parameter Settings

We used pathway2vec [19] to obtain pathway and EC features using “crt” as the embedding method with the following settings: the number of memorized domain was 3, the explore and the in-out hyperparameters were 0.55 and 0.84, respectively, the number of sampled path instances was 100, the walk length was 100, the embedding dimension size was *m* = 128, the neighborhood size was 5, the size of negative examples was 5, and the used configuration of MetaCyc was “uec”, indicating links among ECs were trimmed. The obtained features were used to leverage correlations among ECs and pathways for training leADS (see Section A.4.3). We then trained leADS using the following default settings (unless otherwise mentioned): the learning rate was 0.0001, the batch size was 50, the number of epochs was 3, the number of models was *g* = 3, the proportion of examples (**per**%) to be selected was 30%, the rate of examples to be discarded was *γ* = 0.1, the number of subsampled pathways for each member was 500, and the cutoff threshold for predictions was 0.5. For the regularized hyperparameter *λ*, we performed 10-fold cross-validation on BioCyc T2 &3 data and found the settings *λ* = 10 to be optimum according to results obtained on golden T1 and CAMI datasets.

#### A.4.3 Incorporating enzyme features

We applied the RUST-norm (“crt”) random walk method from pathway2vec [19] that uses a unit-circle equation to obtain enzyme features with settings provided in Section A.4.2. Then, we use enzyme features to concatenate each example *i* according to:

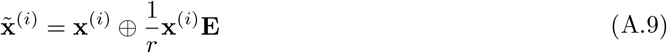

where ⊕ indicates the vector concatenation operation, **E** ∈ℝ^*r*×*m*^ corresponds the feature matrix of enzymes and *m* = 128. The addition of features results in a dimension of size *r* + *m*, where *r* = 3650. We expect by incorporating enzymatic reaction features into the original *r* dimensional example **x**^(*i*)^, the modified 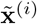 summarizes informative characteristics, which are expected to be useful in pathway prediction.

### A.5 Results and Discussion

#### A.5.1 Parameter Sensitivity

##### Experimental setup

In this section, the impact of two user defined hyperparameters (*k* and **per**%) were evaluated on the CAMI dataset using acquisition functions described in Section A.2.2. In the case of *k*, a range of values between {5, 10, 15, 20, 30, 40, 50, 70, 90, 100} was tested in relation to pathway size for variation ratios in Eq. A.5 or top *k* relevant pathways for nPSP in Eq. A.6. In the case of **per**% different subsampling proportions between {30%, 50%, 70%} were tested by selecting BioCyc T2 &3 data at random. For variation ratios and nPSP, the values of *k* were fixed based on the optimum results obtained from the previous experiment. All other hyperparameters, were set according to the configurations described in Section A.4.2 and results were reported using average F1 scores.

##### Experimental results

Fig. 3a shows the impact of *k* for both variation ratios and nPSP acquisition functions. Although both functions have similar disagreement metrics, the optimum performance for variation ratios is at *k* = 15 while the optimum for nPSP is at *k* = 40. This discrepancy in *k* values likely results from the effects of subsampling pathways and examples that are allocated randomly to each member in *E*. After several rounds of experiments, we found *k* = 50 to be optimum for both variation ratios and nPSP. Next, we examined the effect of **per**% on leADS’s performances using four acquisition functions and random sampling, where we fixed *k* = 50 for variation ratios and nPSP. From Fig. 3b, it is evident that leADS performance generally improves by including more examples for each acquisition function, although the entropy function resulted in a marginal improvement. In contrast, random sampling had no performance benefit across the sample size range tested. In summary, this experiment suggests to consider using **per**% = 70% for any configurations and *k* = 50 for both variation ratios and nPSP.

**Figure 3:**
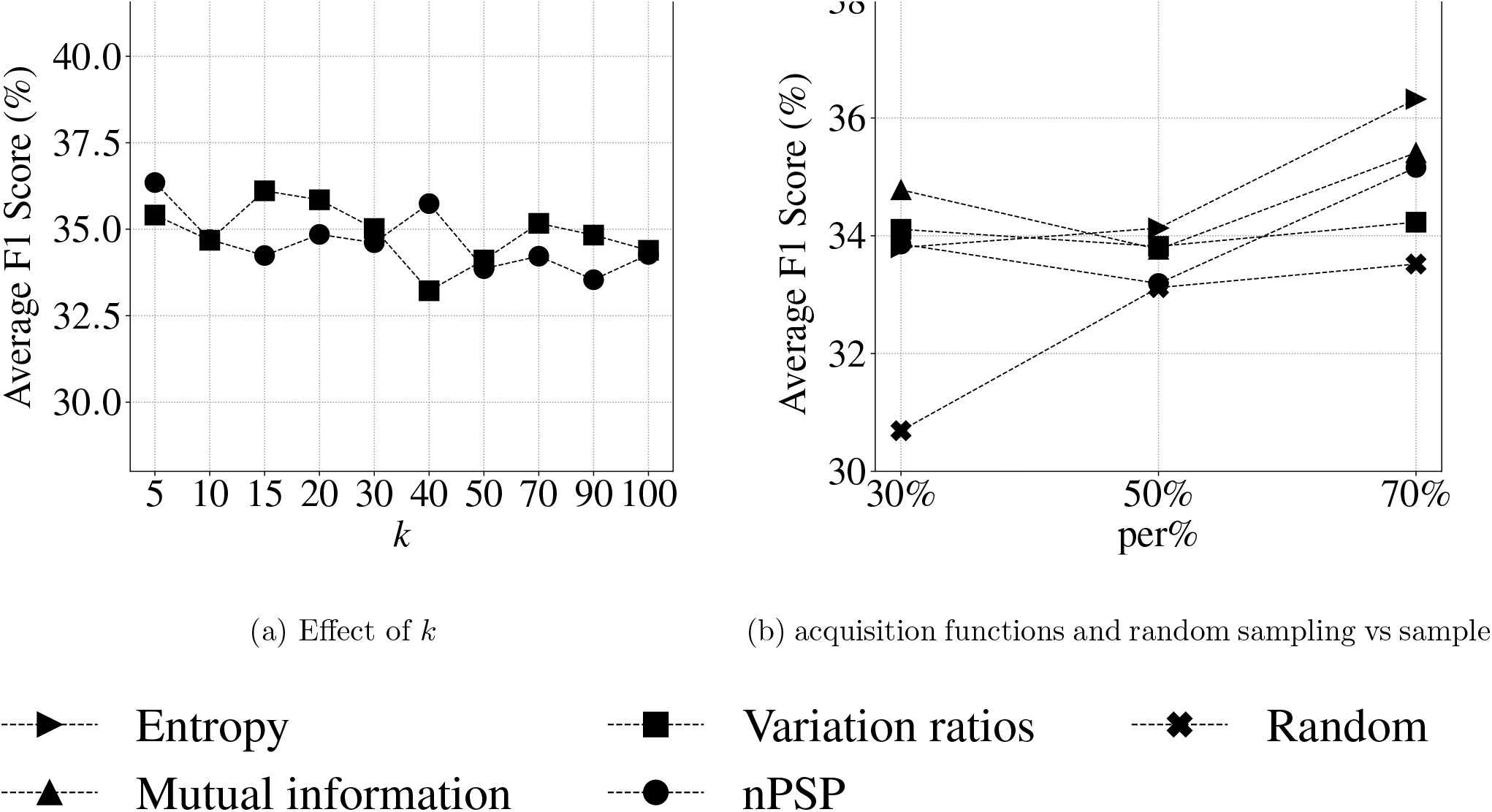
The impact of *k* on leADS performance on the CAMI dataset by varying *k* ∈{5, 10, 15, 20, 30, 40, 50, 70, 90, 100} using variation ratios and nPSP as acquisition functions is demonstrated in Fig. 3a while the effect of four acquisition functions and random sampling by varying sample size according to **per**% ∈ {30%, 50%, 70%} is shown in Fig. 3b.

#### A.5.2 Scalability to the Ensemble Size

##### Experimental setup

In this section, time complexity of training was determined when the model size varied as *g* ∈{1, 2, 3, 5, 10, 15, 20, 50}, simultaneously. Performance was evaluated on the CAMI dataset as described above using the average F1 score metric for each configuration of *g*. **per**% was set to 30% of BioCyc v21 T2 &3 data for training under the four acquisition functions. In the case of random sampling, leADS was trained on 30% of randomly selected BioCyc v21 T2 &3 data. Performance was expected to improve proportionally to the member size in *E* (due to the dual effects of pathways and examples that are being allocated randomly to each base learner) with concomitant increase in computational time. See section A.4.2 for configuration settings.

##### Experimental results

Results in Fig. 4a are consistent with expectations, with gradual inclusion Acquisition functions and random sampling vs model size Model size vs training time of more members in *E* improving leADs performance. Although random sampling observed to have the lowest computational time complexity, nonetheless, this method resulted the lowest performance according to the average F1 metric (Fig. 4b). Among the four acquisition functions, variation ratios required an additional mode operation, contributing to increased training time. Based on these results, we suggest to set the member size between [3, 10] ∈𝕫_*>*0_ while increasing pathway subsampling size accordingly (e.g. 2000 for 10 members) to improve prediction outcomes and reduce both computational complexity (training and inference) and model size.

**Figure 4:**
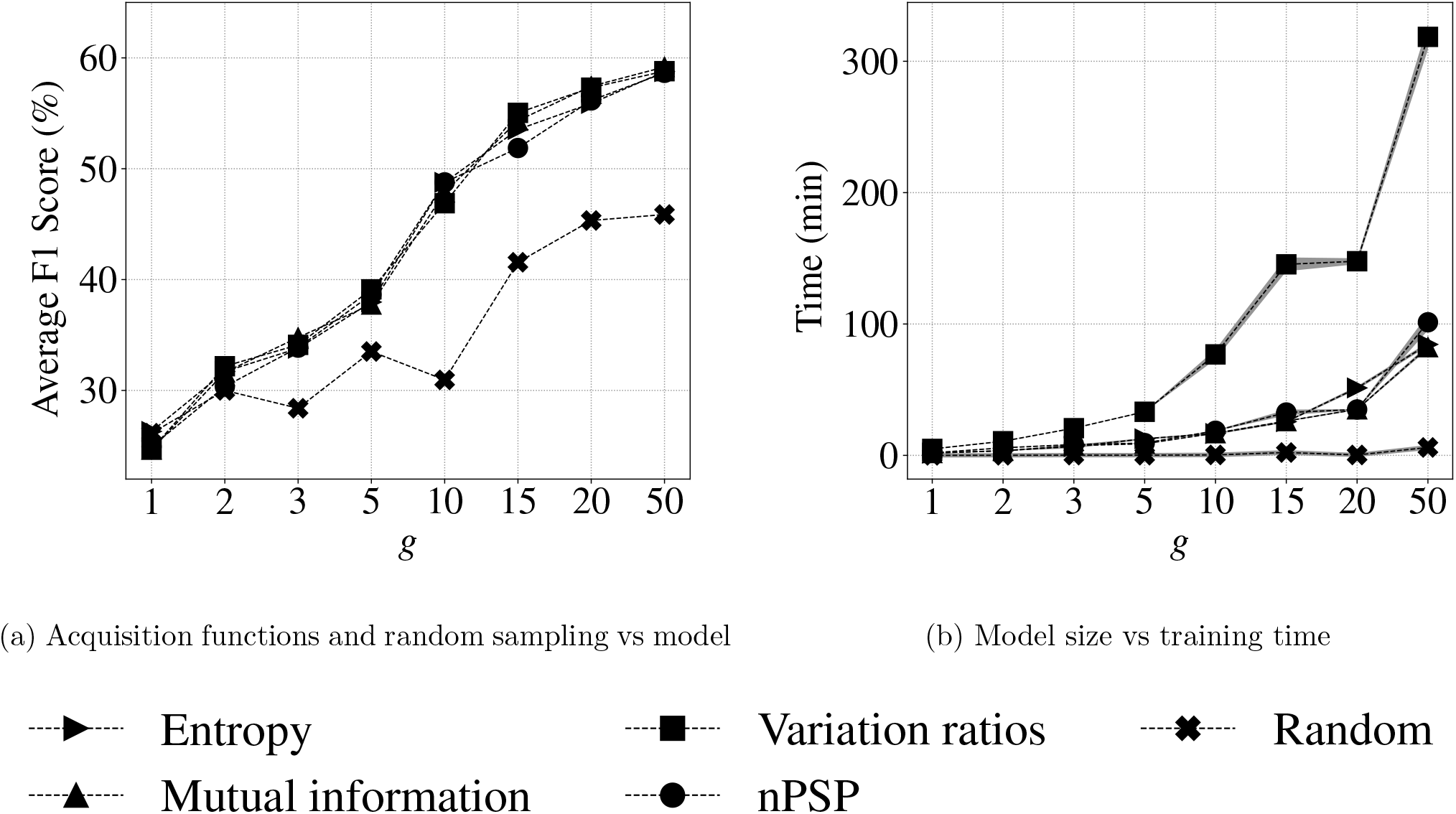
Fig. 4a shows the average F1 score reported on CAMI data as the ensemble size *g* varies across {1, 2, 3, 5, 10, 15, 20, 50} while the elapsed computational time (in minutes) per epoch (averaged over 3 epochs) is demonstrated in Fig. 4b based on the same ensemble size variation.

#### A.5.3 Metabolic Pathway Prediction

##### Experimental setup

In this section, pathway prediction performance was evaluated using parameter settings described in Section A.4.2. Three training configurations were tested: i)- **per**% = 70% under the four acquisition functions, ii)- random sampling corresponding to 70% of BioCyc T2 &3 selected at random, and iii)- full configuration where all BioCyc T2 &3 data were utilized without subsampling. After training, pathway prediction results were reported on golden T1 data using four evaluation metrics: *Hamming loss, average precision, average recall*, and *average F1 score*. leADS performance was compared to three pathway prediction algorithms: i)- MinPath v1.2 [31], ii)- PathoLogic v21 [13]; and iii)- mlLGPR [20] on the T1 data. In addition, we compared leADS performance to other methods on multi-organismal datasets including symbiont, CAMI low complexity and HOTS datasets. For all experiments, the number of epochs was 10, the member size was *g* = 10, the subsampled pathway size was 2000, and *k* was 50 (for variation ratios and nPSP). See Section A.4.2 for additional configuration settings.

##### Experimental results

As shown in Table 2, leADS resulted in competitive performance compared to other pathway inference algorithms based on average F1 scores. For each column in Table 2, a boldface number represents the best evaluation metric score while an underlined number indicates the best score between leADS variants. Among the four acquisition functions, leADS+nPSP resulted in the highest average F1 scores for EcoCyc (0.8874), HumanCyc (0.8333), and TrypanoCyc (0.6897) which are also the highest scores among all models tested. On the other hand, random sampling achieved the poorest overall performance scores. Interestingly, leADS+F in Table 2 was on par with random sampling, reinforcing the idea that BioCyc v21 T2 &3 contain noisy data that hampered proper estimation of leADS coefficients. Through subsampling examples, leADS was able to reduce noise and improve the prediction performance on golden T1 data.

**Table 2:** Predictive performance of each comparing algorithm on 6 benchmark datasets. leADS+F: leADS with full data, leADS+R: leADS with random sampling, leADS+ℋ: leADS with entropy, leADS+_ℳ_: leADS with mutual information, leADS+_𝒱_: leADS with variation ratios, and leADS+nPSP: leADS with normalized propensity scored precision. For each performance metric, ‘↓’ indicates the smaller score is better while ‘↑’ indicates the higher score is better. Values in boldface represent the best performance score while the underlined score indicates the best performance among leADS variances.

| Methods | Hamming Loss ↓ |  |  |  |  |  |
| --- | --- | --- | --- | --- | --- | --- |
|  | EcoCyc | HumanCyc | AraCyc | YeastCyc | LeishCyc | TrypanoCyc |
| PathoLogic | 0.0610 | 0.0633 | 0.1188 | **0.0424** | 0.0368 | **0.0424** |
| MinPath | 0.2257 | 0.2530 | 0.3266 | 0.2482 | 0.1615 | 0.2561 |
| mlLGPR | 0.0804 | 0.0633 | **0.1069** | 0.0550 | 0.0380 | 0.0590 |
| leADS+R | 0.0574 | 0.0796 | 0.1528 | 0.0796 | 0.0515 | 0.0685 |
| leADS+F | 0.0471 | 0.0732 | 0.1576 | 0.0736 | 0.0396 | 0.0566 |
| leADS+$\mathcal{H}$ | 0.0265 | 0.0610 | 0.1453 | 0.0756 | 0.0471 | 0.0606 |
| leADS+$\mathcal{M}$ | 0.0289 | 0.0499 | 0.1425 | 0.0657 | 0.0408 | 0.0542 |
| leADS+$\mathcal{V}$ | 0.0301 | 0.0424 | <u>0.1394</u> | <u>0.0649</u> | 0.0368 | 0.0507 |
| leADS+nPSP | **0.0261** | **0.0364** | 0.1457 | 0.0653 | **0.0333** | <u>0.0499</u> |
| Methods | Average Precision ↑ |  |  |  |  |  |
|  | EcoCyc | HumanCyc | AraCyc | YeastCyc | LeishCyc | TrypanoCyc |
| PathoLogic | 0.7230 | 0.6695 | 0.7011 | **0.7194** | 0.4803 | 0.5480 |
| MinPath | 0.3490 | 0.3004 | 0.3806 | 0.2675 | 0.1758 | 0.2129 |
| mlLGPR | 0.6187 | 0.6686 | 0.7372 | 0.6480 | 0.4731 | 0.5455 |
| leADS+R | 0.7516 | 0.6383 | 0.7199 | 0.5714 | 0.3799 | 0.5039 |
| leADS+F | 0.7994 | 0.6767 | 0.7171 | 0.6352 | 0.4606 | 0.5611 |
| leADS+$\mathcal{H}$ | **0.9380** | 0.6997 | 0.7299 | 0.5872 | 0.4192 | 0.5423 |
| leADS+$\mathcal{M}$ | 0.9239 | 0.7508 | 0.7757 | 0.6684 | 0.4529 | 0.5779 |
| leADS+$\mathcal{V}$ | 0.9231 | 0.7654 | 0.8110 | 0.6720 | 0.4828 | 0.6009 |
| leADS+nPSP | 0.9319 | **0.8425** | **0.8198** | <u>0.7078</u> | **0.5102** | **0.6061** |
| Methods | Average Recall ↑ |  |  |  |  |  |
|  | EcoCyc | HumanCyc | AraCyc | YeastCyc | LeishCyc | TrypanoCyc |
| PathoLogic | 0.8078 | 0.8423 | 0.7176 | 0.8734 | 0.8391 | 0.7829 |
| MinPath | **0.9902** | **0.9713** | **0.9843** | **1.0000** | **1.0000** | **1.0000** |
| mlLGPR | 0.8827 | 0.8459 | 0.7314 | 0.8603 | 0.9080 | 0.8914 |
| leADS+R | 0.7883 | 0.6452 | 0.3980 | 0.4891 | 0.7816 | 0.7429 |
| leADS+F | 0.8176 | 0.6452 | 0.3627 | 0.4410 | 0.8736 | <u>0.8400</u> |
| leADS+$\mathcal{H}$ | 0.8371 | 0.7849 | <u>0.4451</u> | <u>0.5590</u> | 0.9540 | 0.8057 |
| leADS+$\mathcal{M}$ | 0.8306 | 0.8208 | 0.4137 | 0.5459 | 0.8851 | 0.8057 |
| leADS+$\mathcal{V}$ | 0.8208 | <u>0.8889</u> | 0.4039 | 0.5546 | <u>0.9655</u> | 0.8000 |
| leADS+nPSP | <u>0.8469</u> | 0.8244 | 0.3569 | 0.4760 | 0.8621 | 0.8000 |
| Methods | Average F1 ↑ |  |  |  |  |  |
|  | EcoCyc | HumanCyc | AraCyc | YeastCyc | LeishCyc | TrypanoCyc |
| PathoLogic | 0.7631 | 0.7460 | 0.7093 | **0.7890** | 0.6109 | 0.6447 |
| MinPath | 0.5161 | 0.4589 | 0.5489 | 0.4221 | 0.2990 | 0.3511 |
| mlLGPR | 0.7275 | 0.7468 | **0.7343** | 0.7392 | 0.6220 | 0.6768 |
| leADS+R | 0.7695 | 0.6417 | 0.5126 | 0.5271 | 0.5113 | 0.6005 |
| leADS+F | 0.8084 | 0.6606 | 0.4818 | 0.5206 | 0.6032 | 0.6728 |
| leADS+$\mathcal{H}$ | 0.8847 | 0.7399 | <u>0.5530</u> | 0.5727 | 0.5825 | 0.6483 |
| leADS+$\mathcal{M}$ | 0.8748 | 0.7842 | 0.5396 | 0.6010 | 0.5992 | 0.6730 |
| leADS+$\mathcal{V}$ | 0.8690 | 0.8226 | 0.5393 | <u>0.6077</u> | **0.6437** | 0.6863 |
| leADS+nPSP | **0.8874** | **0.8333** | 0.4973 | 0.5692 | 0.6410 | **0.6897** |

Metabolic interactions are integral to microbial community structure and function. In some cases these interactions are related to production of public goods by a subset of community members that provision non-producing members, or through the removal of end products enabling unfavorable reactions to proceed [8]. In other cases enzymatic steps within a multi-step pathway are distributed between multiple community members resulting in emergent metabolic properties that are robust to loss of individual members [17]. To evaluate leADS performance on metabolic pathways distributed between organisms we used the reduced genomes of mealybug symbionts *Moranella* (GenBank NC- 015735) and *Tremblaya* (GenBank NC-015736) [21]. Fig. 5A illustrates the distributed genes for the *Lysine biosynthesis* pathway. The two symbiont genomes in combination encode intact biosynthetic pathways for 9 essential amino acids (Fig. 5B where a grey circle corresponds the missing gene by MetaPathways software [15]). PathoLogic, mlLGPR, and leADS were used to predict pathways on individual symbiont genomes and a concatenated dataset consisting of both symbiont genomes, and resulting amino acid biosynthetic pathway distributions were determined (Fig. 5C). PathoLogic and leADS predicted 6 of the expected amino acid biosynthetic pathways on the composite genome while mlLGPR predicted 8 pathways. The *L-phenylalanine biosynthesis I* pathway was not inferred because the associated genes were reported to be missing during the ORF prediction process while *Chorismate biosynthesis I* was not incorporated during training (Fig. 6). All models inferred false positive pathways for individual symbiont genomes (*Moranella* and *Tremblaya*) despite reduced pathway coverage information (mapping enzymes onto associated 9 amino acid biosynthetic pathways) relative to the composite genome (Fig. 7). Although it is possible for leADS to reduce type I error by incorporating taxonomy-based predictions using rules, such pruning can also increase false-negative (type II error) pathway predictions in multi-organismal datasets [9].

**Figure 5:**
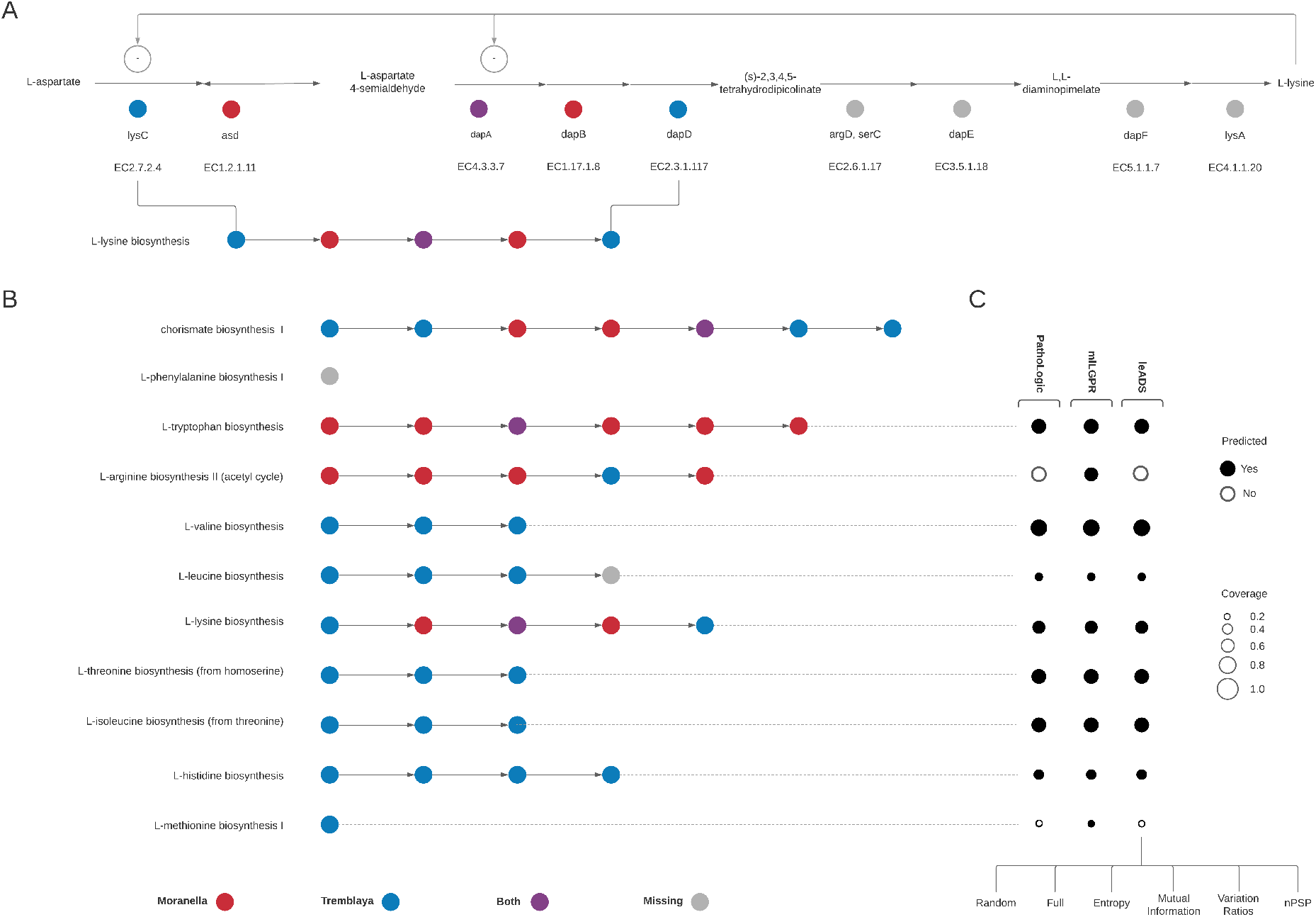
Schematic representation or distributed amino acid biosynthesis in the mealybug symbiotic system. A) indicates a simplified version of the L-lysine biosynthetic pathway from MetaCyc including locus ids and enzyme commission (EC) numbers for each step in the pathway starting from L-aspartate conversion. The specific steps encoded by Moranella (red), Tremblaya (blue) or both (purple) endosymbionts are shown as coloured circles in a simplified glyph structure based on McCuthcheon [21]. Missing steps are indicated in grey. B) Breakdown of essential amino acid biosynthetic pathways in the mealybug symbiosis using simplified glyph structures. C) Corresponding pathway prediction outcomes using PathoLogic, mlLGPR, and leADS (with random sampling, full data, and four acquisition functions). Black circles indicate predicted pathways by associated models while open circles indicate pathways that were not recovered by models. The size of circles corresponds to pathway coverage information used in metabolic inference.

**Figure 6:**
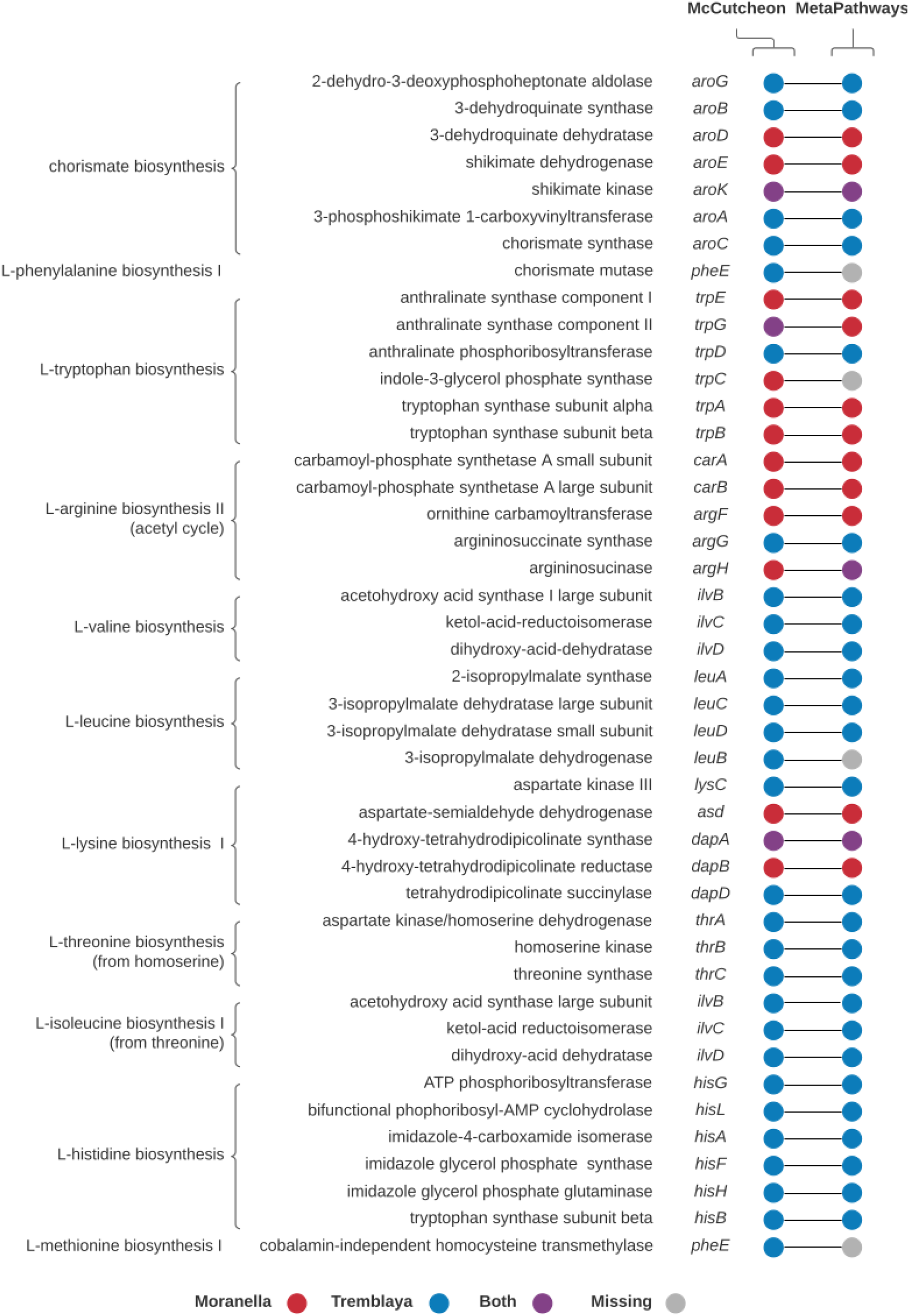
Comparison between Mccutcheon and colleagues [21] reported distributed genes in 9 amino acid pathways in the *Candidatus Moranella endobia* and *Candidatus Tremblaya princeps* genomes with MetaPathways v2.5 [15]. Circles represent detected presence of the pathway enzymes in Moranella (red), Tremblaya (blue), both genomes (purple), or missing (grey).

**Figure 7:**
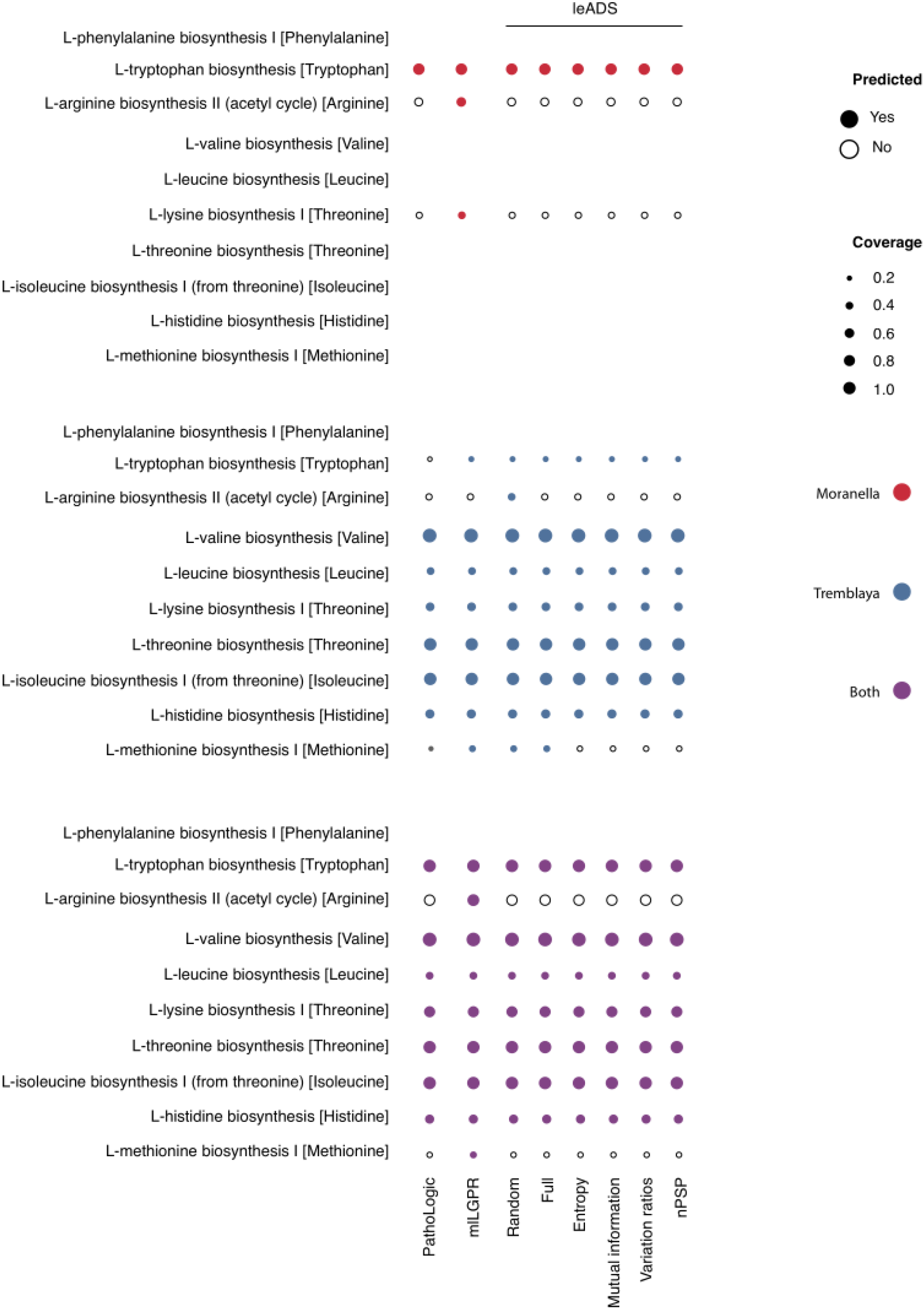
Comparative study of predicted pathways for symbiont data between PathoLogic, mlLGPR, and leADS (with random sampling, full data, and four acquisition functions). Filled red, blue, and purple circles represent detected presence of the pathway in Moranella, Tremblaya, and both genomes, respectively. The size of circles corresponds the pathway coverage information.

To evaluate performance on more complex multi-organismal genomes we compared leADS to mlL-GPR using the CAMI low complexity dataset [24] and to PathoLogic and mlLGPR using the HOTS dataset [29]. In the case of CAMI, leADS+nPSP outperformed other methods resulting in an average F1 score of 0.6214 (Table 3). In the case of HOTS, leADS+R, leADS+F, leADS+ℋ, leADS+ ℳ, leADS+ 𝒱, and leADS+nPSP predicted a total of 60, 67, 63, 68, 67, and 68 pathways among a subset of 180 selected water column pathways [9], while PathoLogic and mlLGPR inferred 54 and 62 pathways, respectively (Figs 8, 9, 10, and 11). These observations indicate that leADS with subsampling improves pathway prediction outcomes by reducing training loss due to the class-imbalance problem in BioCyc v21 data. Based on these results we recommend using nPSP with *g* = 10 and *k* = 50 settings for optimal leADS performance.

**Table 3:** Predictive performance of mlLGPR and leADS on CAMI low complexity data. leADS+F: leADS with full data, leADS+R: leADS with random sampling, leADS+ℋ: leADS with entropy, leADS+ℳ: leADS with mutual information, leADS+𝒱: leADS with variation ratios, and leADS+nPSP: leADS with normalized propensity scored precision. Values in boldface represent the best performance score while the underlined score indicates the best performance among leADS variances.

| Metric | mLGPR | leADS+R | leADS+F | leADS+$\mathcal{H}$ | leADS+$\mathcal{M}$ | leADS+$\mathcal{V}$ | leADS+nPSP |
| --- | --- | --- | --- | --- | --- | --- | --- |
| Hamming Loss (↓) | 0.0975 | 0.0577 | 0.0553 | 0.0402 | 0.0398 | 0.0399 | **<u>0.0397</u>** |
| Average Precision (↑) | 0.3570 | 0.5245 | 0.5468 | 0.7515 | 0.7558 | 0.7550 | **<u>0.7569</u>** |
| Average Recall (↑) | **0.7827** | 0.5212 | 0.5284 | 0.5260 | 0.5306 | 0.5268 | <u>0.5334</u> |
| Average F1 (↑) | 0.4866 | 0.5174 | 0.5320 | 0.6151 | 0.6199 | 0.6167 | **<u>0.6214</u>** |

**Figure 8:**
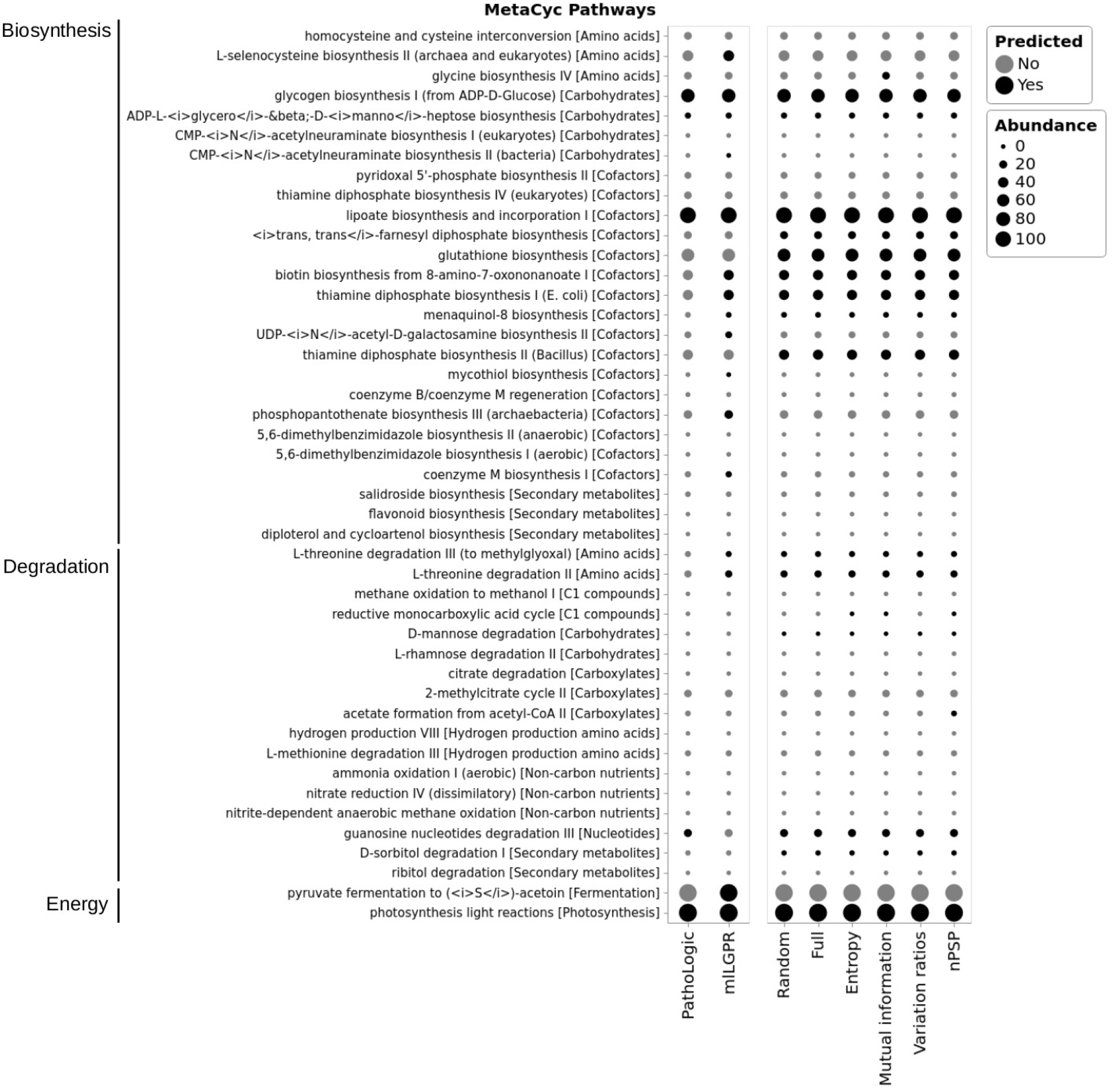
Comparative study of predicted pathways for HOTS 25m dataset between PathoLogic, mlLGPR, and leADS (with random sampling, full data, and four acquisition functions). Black circles indicate predicted pathways by the associated models while grey circles indicate pathways that were not recovered by models. The size of circles corresponds the pathway abundance information.

**Figure 9:**
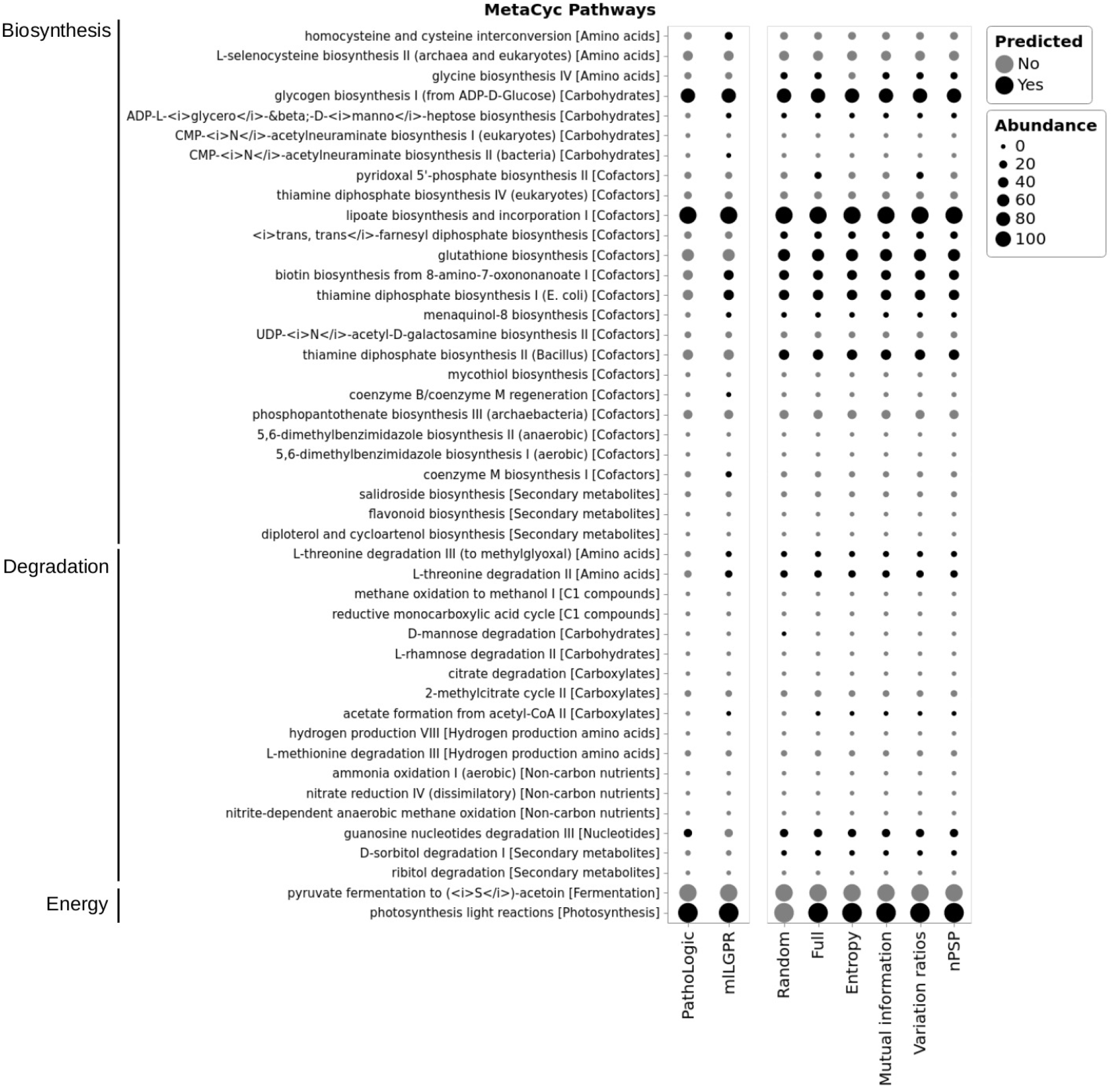
Comparative study of predicted pathways for HOTS 75m dataset between PathoLogic, mlLGPR, and leADS (with random sampling, full data, and four acquisition functions). Black circles indicate predicted pathways by the associated models while grey circles indicate pathways that were not recovered by models. The size of circles corresponds the pathway abundance information.

**Figure 10:**
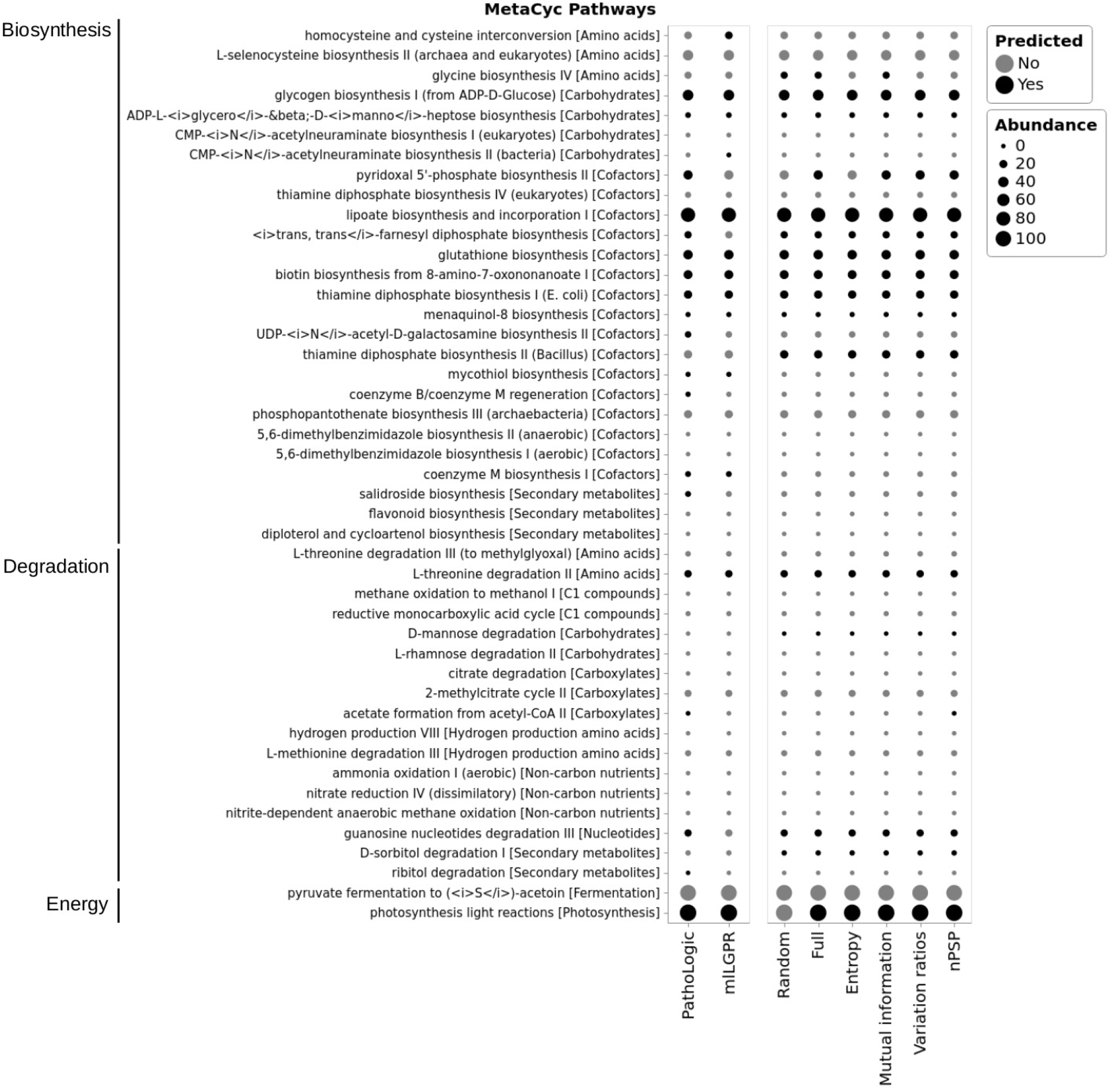
Comparative study of predicted pathways for HOTS 110m dataset between PathoLogic, mlL-GPR, and leADS (with random sampling, full data, and four acquisition functions). Black circles indicate predicted pathways by the associated models while grey circles indicate pathways that were not recovered by models. The size of circles corresponds the pathway abundance information.

**Figure 11:**
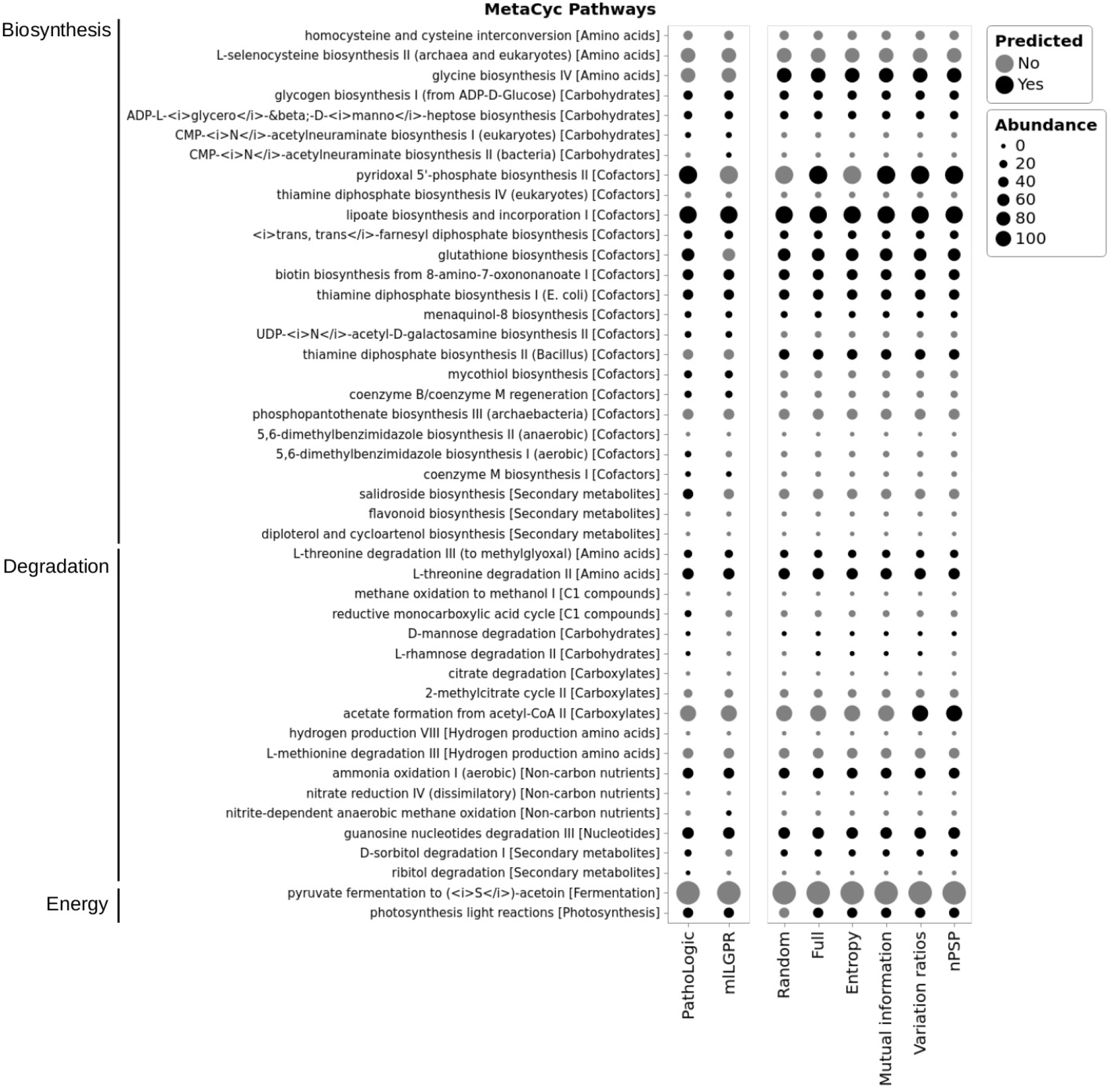
Comparative study of predicted pathways for HOTS 500m dataset between PathoLogic, mlL-GPR, and leADS (with random sampling, full data, and four acquisition functions). Black circles indicate predicted pathways by the associated models while grey circles indicate pathways that were not recovered by models. The size of circles corresponds the pathway abundance information.

## Availability of Data and Materials

The leADS source code is available under the GNU License at github.com/hallamlab/leADS. A wiki, including a tutorial, is available at github.com//hallamlab/leADS/wiki.

## Acknowledgments

We would like to thank Connor Morgan-Lang, Kishori Konwar, and Aria Hahn for lucid discussions on the function of the leADS model and all members of the Hallam Lab for helpful comments along the way.

